# Associations of Socioeconomic Disparities With Buccal DNA-Methylation Measures Of Biological Aging

**DOI:** 10.1101/2022.12.07.519438

**Authors:** L. Raffington, T. Schwaba, M. Aikins, D. Richter, G.G. Wagner, K.P. Harden, D.W. Belsky, E.M. Tucker-Drob

## Abstract

**Background:** Individuals who are socioeconomically disadvantaged are at increased risk for aging-related diseases and perform less well on tests of cognitive function. The Weathering Hypothesis proposes that these disparities in physical and cognitive health arise from an acceleration of biological processes of aging. Theories of how life adversity is biologically embedded identify epigenetic alterations, including DNA methylation (DNAm), as a mechanistic interface between the environment and health. Consistent with the Weathering hypothesis and theories of biological embedding, recently developed DNAm algorithms have revealed profiles reflective of more advanced aging and lower cognitive function among socioeconomically-at-risk groups. These DNAm algorithms were developed using blood-DNA, but social and behavioral science research commonly collect saliva or cheek-swab DNA. This discrepancy is a potential barrier to research to elucidate mechanisms through which socioeconomic disadvantage affects aging and cognition. We therefore tested if social gradients observed in blood-DNAm measures could be reproduced using buccal-cell DNA obtained from cheek swabs.

**Results:** We analyzed three DNAm measures of biological aging and one DNAm measure of cognitive performance, all of which showed socioeconomic gradients in previous studies: the PhenoAge and GrimAge DNAm clocks, DunedinPACE, and Epigenetic-*g*. We first computed blood-buccal cross-tissue correlations in n=21 adults (GEO111165). Cross-tissue correlations were low-to-moderate across (*r*=.25 to *r*=.48). We next conducted analyses of socioeconomic gradients using buccal DNAm data from SOEP-G (n=1128, 57% female; age mean=42 yrs, SD=21.56, range 0-72). Associations of socioeconomic status with DNAm measures of aging were in the expected direction, but were smaller as compared to reports from blood DNAm datasets (*r*=-.08 to *r*=-.13).

**Conclusions:** Our findings are consistent with the hypothesis that socioeconomic disadvantage is associated with DNAm indicators of worse physical and cognitive health. However, relatively low cross-tissue correlations and attenuated effect-sizes for socioeconomic gradients in buccal DNAm compared with reports from analysis of blood DNAm suggest that, in order to take full advantage of buccal-DNA samples, DNAm algorithms customized to buccal DNAm are needed.

## Background

Individuals who are socioeconomically disadvantaged are at increased risk for aging-related diseases and exhibit lower average levels of cognitive function across the life course, (Gkiouleka et al., 2018; Lövdén et al., 2020, p. 202; Tucker-Drob, 2019). Studies of humans and other animals identify several biological pathways through which social factors drive disease, including dysregulation of immune and metabolic systems in response to chronic stress (Snyder-Mackler et al., 2020). These pathways overlap substantially with the biology that mediates aging-related health declines (López-Otín et al., 2013). This overlap is consistent with the Weathering Hypothesis, which proposes that social adversity accelerates biological processes of aging (Geronimus et al., 2006).

Biological aging can be conceptualized as the progressive loss of system integrity that occurs with advancing chronological age (Kirkwood, 2005). The current state-of-the-art for quantification of biological aging in epidemiological studies of humans is a family of DNA methylation (DNAm) measurements. Epigenetic changes, including DNAm, are among the hallmarks of aging and are theorized to be key transducers of biological embedding of social adversity (Hertzman & Boyce, 2010; López-Otín et al., 2013). DNAm measures of biological aging that are most strongly predictive of disease, disability, and mortality are also consistently associated with social determinants of health (Oblak et al., 2021; Raffington & Belsky, 2022). In addition, there is evidence for social patterning of a DNAm measurement quantifying cognitive performance (McCartney et al., 2022), which parallels well-documented socioeconomic disparities in cognitive function across the life course (Lövdén et al., 2020). These DNAm measures open opportunities to study mechanisms of social disparities in physical and cognitive health and to guide the development and evaluation of interventions to address them.

A barrier to achieving this potential is that DNAm is specific to types of tissues and cells; it is a critical mechanism of cellular differentiation and determinant of cellular phenotype (Bakulski et al., 2016). Most DNAm algorithms used to study social gradients in health were developed from analysis of DNA derived from blood samples. Therefore, the ideal setting for their application is blood-derived DNA methylation. However, collection of blood samples is not feasible in some studies. For these studies, alternative sources of DNA, such as saliva and buccal tissue (*i*.*e*., inner cheek) may be easier to obtain. The extent to which algorithms developed from blood-derived DNA can provide reliable and valid measurements in alternative tissues remains uncertain.

In two prior projects, we followed up algorithms developed to measure biological aging and cognitive functioning from blood DNAm in saliva samples collected from a pediatric cohort (Raffington, Belsky, et al., 2021; Raffington, Tanksley, et al., in press). In those studies, we were able to replicate several observations made from blood samples. First, the DNAm measure of the pace of biological aging (*i*.*e*., a previous iteration of DunedinPACE) exhibited a parallel socioeconomic gradient in the pediatric saliva samples as had been observed previously in blood DNAm datasets from adults. Second, the DNAm measure of cognitive functioning Epigenetic-*g* exhibited parallel association with children’s performance on cognitive tests as had been observed previously in a blood DNAm dataset from adults. In contrast, the PhenoAge and GrimAge DNAm measures of biological age showed no social gradient in the pediatric saliva samples, in contrast to results from studies of blood samples (Schmitz et al., 2021).

Saliva is composed of a mix of leukocytes (which are also the source of blood-derived DNA samples) and epithelial cells. Buccal sample-derived DNA comes predominantly from epithelial cells. It is unclear whether DNAm measures computed in buccal DNAm will show similar evidence of trans-tissue validation. Here, we examined whether the same socioeconomic gradients in biological aging and DNAm-predicted cognitive performance apparent in blood DNAm analyses could be reproduced in analysis of buccal DNAm. The analysis we report is based on a pre-registration plan filed with OSF (https://osf.io/msjgc). Where our work has developed beyond this original pre-registration, we note it in the text. We first tested cross-tissue correlations of DNAm measures of biological aging (*i*.*e*., PhenoAge Accel., GrimAge Accel., DunedinPACE) and DNAm-predicted cognitive performance (*i*.*e*., Epigenetic-*g*) in buccal and blood DNAm datasets generated from the same individuals using the public dataset GEO111165 (n=21). Next, we examined association of chronological age with buccal DNAm measures in n=1128 participants from SOEP-G (57% female; age mean=42 yrs, SD=21.56, range 0-72). Finally, we tested associations of socioeconomic status with DNAm algorithms computed from buccal-cell DNAm in the same SOEP-G sample.

## Results

### (1) Cross-tissue correlations between blood and buccal samples were low-to-moderate

We evaluated the correspondence between buccal and blood DNAm measures in an auxiliary dataset that collected both buccal and blood samples from the same n=21 people (Braun et al., 2019); Illumina EPIC array dataset in Gene Expression Omnibus accession GSE11116, https://www.ncbi.nlm.nih.gov/geo/query/acc.cgi?acc=GSE111165).

Cross-tissue correlations between blood and buccal samples of the DNAm measures were low-to-moderate across measures (*r*=0.25 to *r*=0.48). Means of DNAm measures were higher in buccal compared to blood samples, with the exception of Epigenetic-*g*, for which mean comparisons are not possible because beta-methylation values are standardized prior to computation (see Table 1).

**Table 1.**
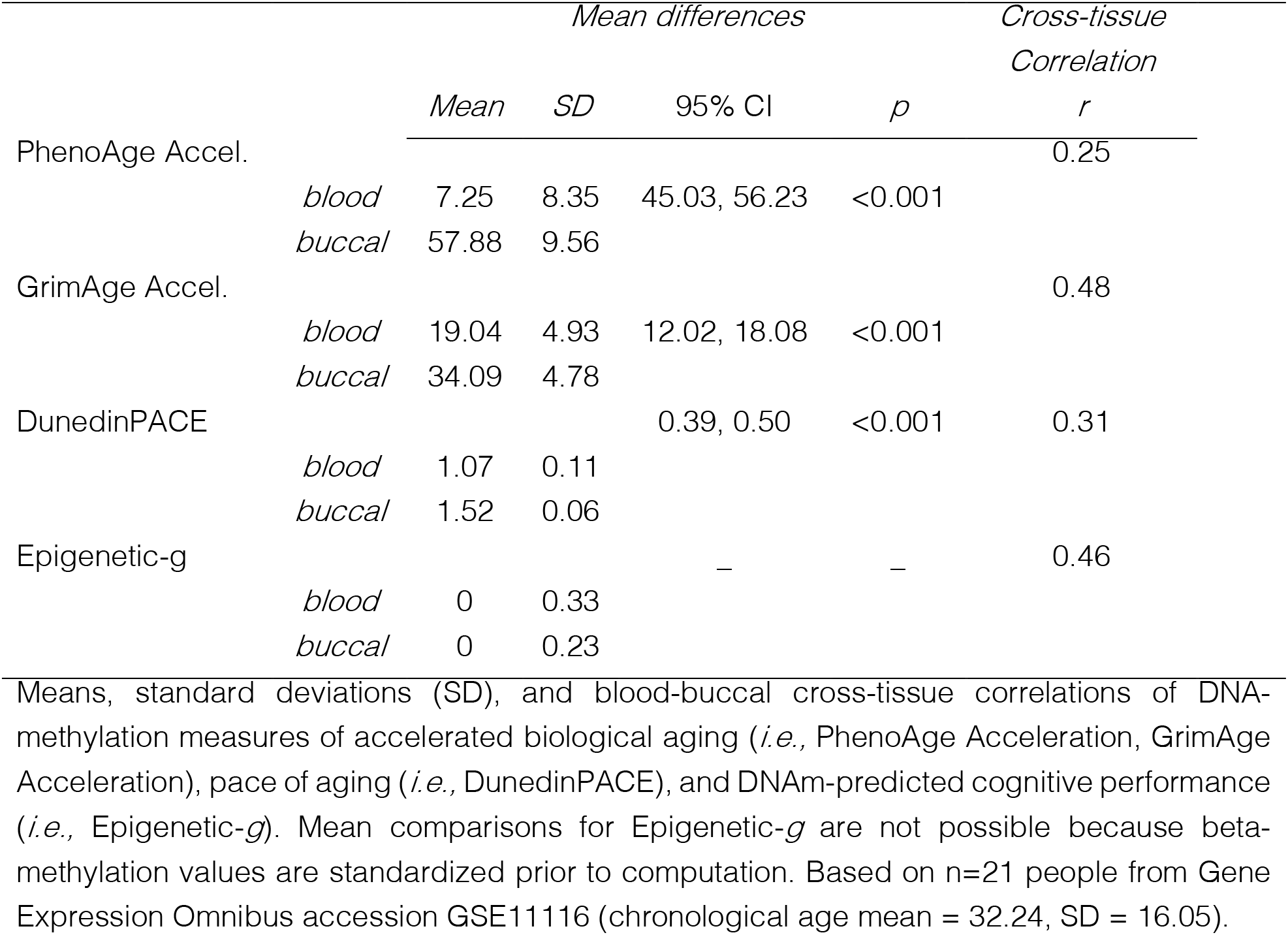
Blood-buccal cross-tissue correlations of blood-based DNA-methylation measures (n=21).

### (2) Chronological age gradients in biological aging are reproduced in buccal DNAm

We examined associations of chronological age with buccal DNAm algorithms. For PhenoAge, strong association with chronological age is expected. In SOEP-G, participants’ buccal DNAm PhenoAge values were highly correlated with their chronological ages (PhenoAge *r*=0.89, 95% CI = 0.88, 0.90, *p*<0.001). GrimAge calculations include information about participant chronological age and, as a result, show very strong correlations (*r*=0.99, 95% CI = 0.99, 0.99, *p*<0.001). In contrast to PhenoAge and GrimAge, which estimate biological age values, DunedinPACE estimates the pace of aging. Consistent with prior reports from blood DNAm datasets and with biodemography theory, which proposes that the pace of aging accelerates as we grow older (Belsky et al., 2022; Finch & Crimmins, 2016), participants’ DunedinPACE values were moderately correlated with their chronological ages (*r* =0.24, 95% CI = 0.18, 0.29, *p*<0.001). We also observed positive age trends for Epigenetic-*g*, mirroring known patterns of cognitive development; values increased across the first half of the lifespan and then stabilized in late middle age (*r* =0.45, 95% CI = 0.40, 0.49, *p*<0.001; age in years unstandardized *b* = 0.008, 95% CI = 0.006 – 0.011, *p*<0.001; age squared unstandardized *b* = -0.001, 95% CI = -0.001-0.000, *p*=0.001). Age patterning of DNAm measures is shown in in Figure 1.

**Figure 1.**
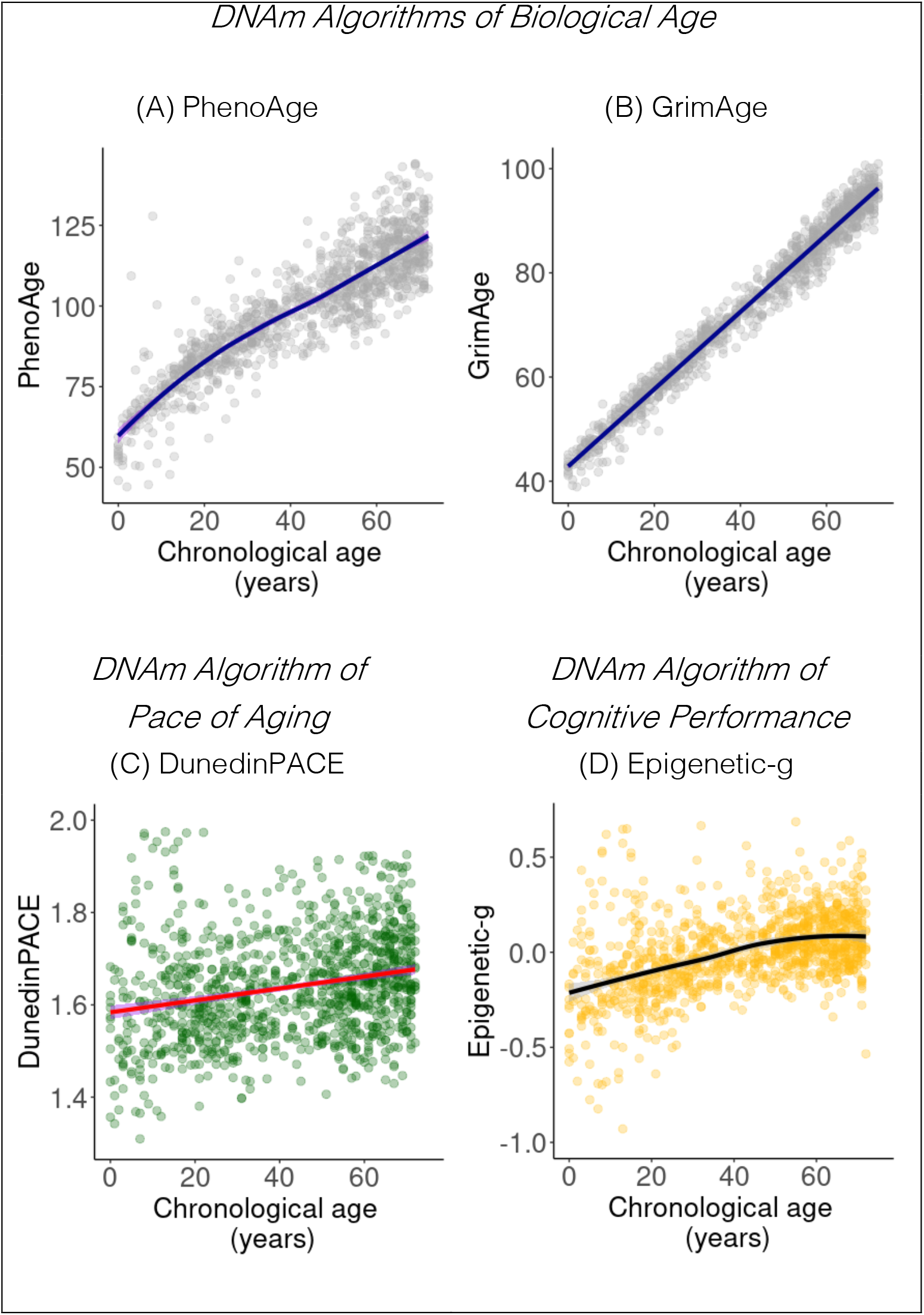
Chronological age and buccal DNAm algorithms. Panel (A-B) plot associations of chronological age with buccal DNAm algorithms of biological aging, for which strong associations are expected: (A) PhenoAge and (B) GrimAge. Panel (C) plots association of chronological age with the pace of aging, DunedinPACE. Panel (D) plots association of chronological age with a DNAm algorithm of cognitive performance, Epigenetic-*g*

### (3) Socioeconomic disadvantage is associated with accelerated biological aging in Germany

We tested associations of socioeconomic status (SES) with DNAm measures of biological aging computed from buccal-cell DNAm in SOEP-G. SES was measured as a composite of household income and educational levels (highest in household). Consistent with reports from blood DNAm datasets, participants with higher SES had younger biological ages and slower pace of aging (*r’s* = -0.08 to -0.13, *p*’s < 0.011, **Table 2**).

**Table 2.**
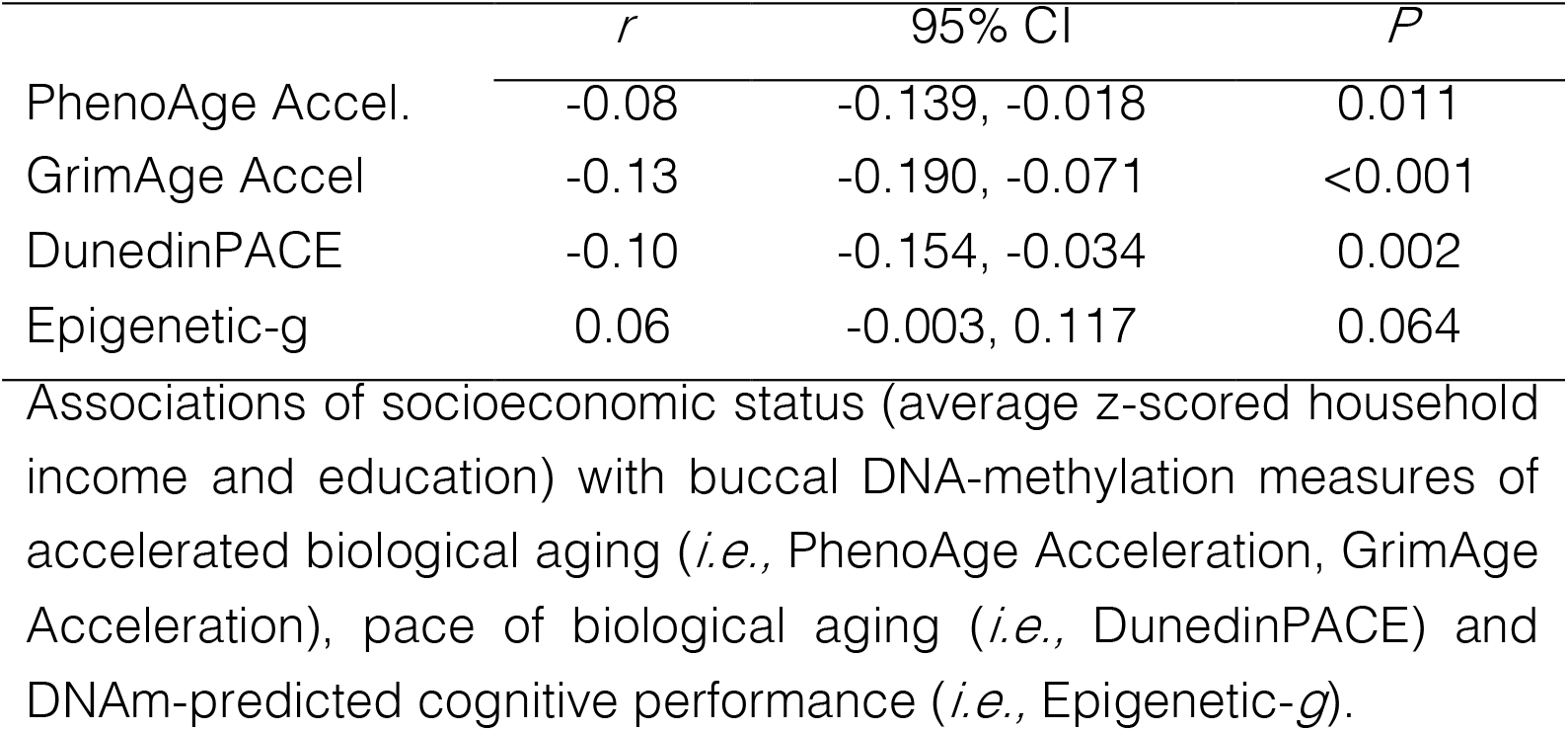
Associations of socioeconomic status with buccal DNA-methylation measures.

|  | *r* | 95% CI | *P* |
| --- | --- | --- | --- |
| PhenoAge Accel. | -0.08 | -0.139, -0.018 | 0.011 |
| GrimAge Accel | -0.13 | -0.190, -0.071 | <0.001 |
| DunedinPACE | -0.10 | -0.154, -0.034 | 0.002 |
| Epigenetic-g | 0.06 | -0.003, 0.117 | 0.064 |

Associations of socioeconomic status (average z-scored household income and education) with buccal DNA-methylation measures of accelerated biological aging (*i.e.*, PhenoAge Acceleration, GrimAge Acceleration), pace of biological aging (*i.e.*, DunedinPACE) and DNAm-predicted cognitive performance (*i.e.*, Epigenetic-*g*).

Next, according to our pre-registered analysis plan, we tested whether the association of SES with DNAm measures of aging differed by chronological age. This interaction was statistically significant for PhenoAge and GrimAge Acceleration (SES by continuous age interaction on PhenoAge std b= -0.11, 95% CI = -0.17, -0.05, *p*<0.001; Grimage std b= -0.07, 95% CI = -0.13, -0.02, *p*=0.011). There were no age differences in the SES association with DunedinPACE (*p*-value for continuous age interaction = 0.916). To further illustrate the interaction, we stratified the sample into older and younger participants. Among the older participants (aged>42 years, n=576), the SES association with PhenoAge Acceleration was *r*= - 0.14, 95% CI = -0.22, -0.06, *p*<0.001 and with GrimAge Acceleration was *r*= -0.18, 95% CI = - 0.26, -0.10, *p*<0.001. In contrast, among younger participants (aged<42 years, n=482), the SES association with PhenoAge Acceleration was *r*= 0.03, 95% CI = -0.06, 0.12, *p*= 0.494 and with GrimAge Acceleration was *r*= -0.04, 95% CI = -0.13, 0.05, *p*=0.352. In sum, SES was associated with PhenoAge and GrimAge Acceleration only for older participants, whereas low SES was associated with DunedinPACE across age groups. **Figure 2** shows the association of socioeconomic status with DNAm by age.

**Figure 2.**
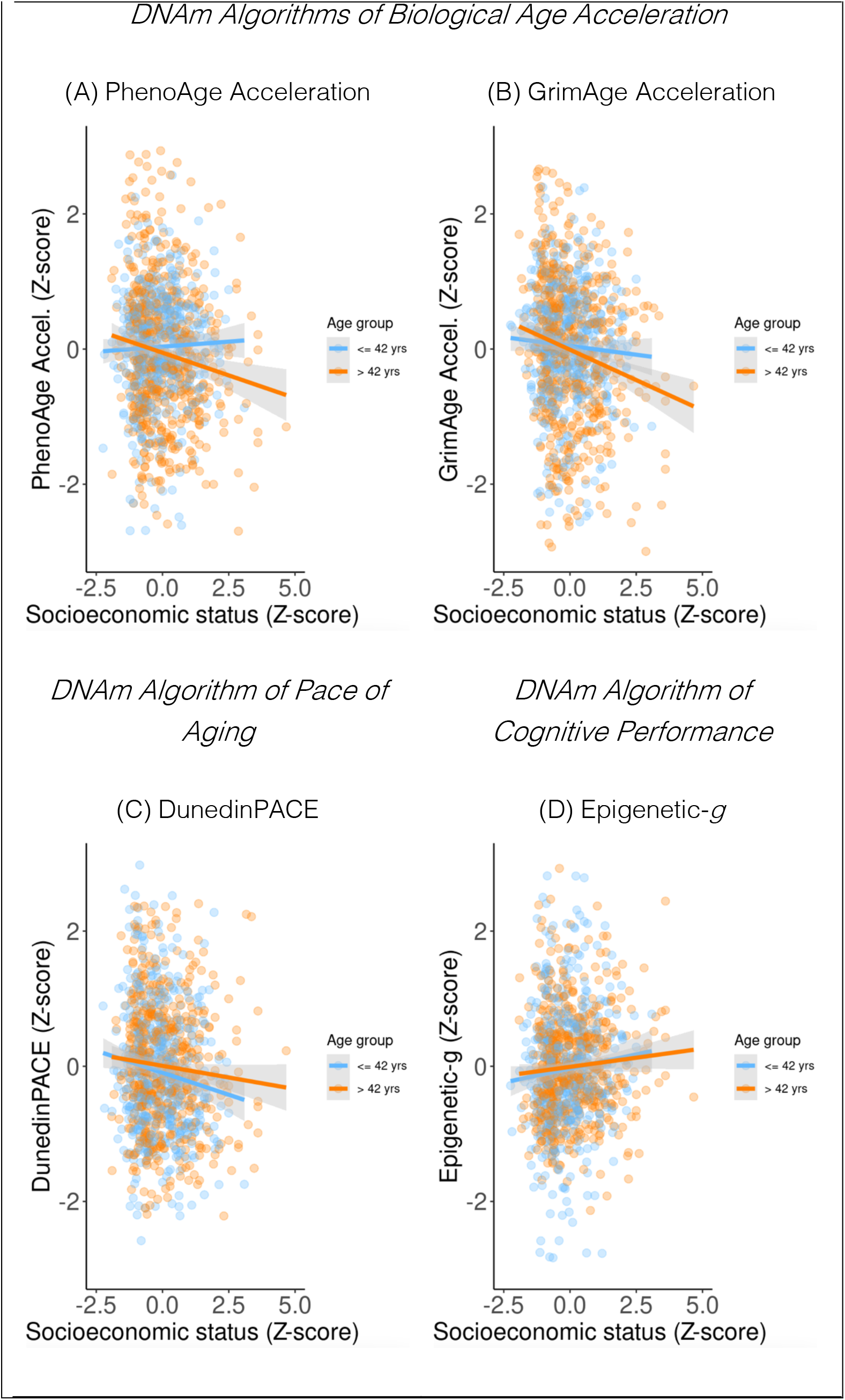
Socioeconomic status and buccal DNAm algorithms. Panel (A-B) plot associations of socioeconomic status with buccal DNAm algorithms of accelerated biological aging: (A) PhenoAge Acceleration and (B) GrimAge Acceleration. Panel (C) plots association of socioeconomic status with the pace of aging, DunedinPACE. Panel (D) plots association of socioeconomic status with a DNAm algorithm of cognitive performance, Epigenetic-*g*

Association of socioeconomic status with Epigenetic-*g* was in the expected direction, but was small and not statistically different from zero at the alpha=0.05 level (see **Table 4** and **Figure 2D**). Excluding smokers and accounting for body mass index did not substantially affect associations with SES (see **Figure S1**).

## Discussion

We tested if socioeconomic gradients in DNAm measurements of biological aging and cognitive performance, which are apparent in blood DNAm analyses, could be reproduced in analysis of buccal DNAm. Our findings are consistent with the Weathering Hypothesis that socioeconomic disadvantage is associated with accelerated biological aging. However, effect-sizes were approximately 50% lower than those reported in previously published analyses of blood DNAm datasets. Such studies have reported associations of magnitude of approximately *r*=.20, ranging from *r*=.10 to *r*=.37 (Raffington & Belsky, 2022), whereas here we report associations of magnitude of approximately *r*=.10, ranging from *r*=.079 to *r*=.13 Similarly, associations of socioeconomic status with buccal DNAm-predicted cognitive performance were attenuated by approximately 50% and not statistically different from zero, in contrast to studies of blood and saliva DNAm datasets, which have reported associations with socioeconomic measures of magnitude *r*=.11 and *r*=.14 (McCartney et al., 2022; Raffington, Tanksley, et al., in press, note larger effect sizes for neighborhood-level socioeconomic contexts). Moreover, cross-tissue correspondence of DNAm indices was low-to-moderate. Collectively, these findings suggest that in order to take full advantage of buccal DNA samples, it will be important to develop DNAm indices that are customized to buccal DNAm.

One observation from our buccal DNAm data is that SES was associated with more PhenoAge Acceleration and GrimAge Acceleration only for older participants, whereas in the case of DunedinPACE the socioeconomic gradient was evident for both young and old participants. This pattern of results is consistent with findings from saliva DNAm in children and adolescents, which showed no association of PhenoAge and GrimAge with household SES, but did identify an association with DunedinPACE (Raffington, Belsky, et al., 2021). One possible explanation for this result is that measures of biological age, such as PhenoAge and GrimAge, which were designed to quantify differences in mortality risk among midlife and older adults, may be less sensitive to early stages in the biological embedding of social disadvantage. Replication of this result in other datasets and across tissues is needed.

## Conclusion

Our findings are consistent with the hypothesis that socioeconomic disadvantage is associated with accelerated biological aging in Germany. However, cross-tissue correspondence of DNAm indices was low-to-moderate and effect-sizes for SES associations estimated from buccal DNAm were attenuated by roughly 50% compared with reports from blood DNAm datasets. Development of DNAm measures of biological aging and cognitive performance that are customized to buccal DNAm should be a research priority.

## Methods

### 1. Participants

SOEP-G participants were from the SOEP-IS cohort, which is based on a random sample of German households and contains a rich array of information on socioeconomic context, household dynamics, personality, and health (Koellinger et al., 2021). 6,576 people were originally invited to participate in the 2019 wave of the SOEP-IS with the aim to collect saliva for genotyping, 2598 of whom provided a valid genetic sample. ∼98% of the genotyped SOEP-IS sample is of high genetic similarity to European reference groups. See Koellinger et al. (2021) for more information on the genotyped SOEP-IS cohort called SOEP-G.

Residual frozen DNA samples from n=1128 individuals from the n=2598 genotyped SOEP-G cohort were selected for DNAm extraction based on the availability of funds (see Table 3 for descriptive statistics). Exclusion and inclusion criteria were: (1) exclusion of 5 samples due to sex mismatch between self-reported and genetic sex, (2) inclusion of all samples from children and adolescents (*i*.*e*., under or equal to 18 yrs) whose residual DNA samples contained at least 50ng of DNA, (3) inclusion of adults that had (a) at least 250ng of DNA left, (b) had a DNA call rate of at least 0.975, (c) were not parents of selected children and adolescents so that the maximum number of different households were included, and (d) extended the age distribution continuously past 18 years so that all younger adults were included. The ID list was randomized so that plate effects were not confounded with chronological age. In addition, 24 samples were randomly selected as technical duplicates. The final sample of n=1128 unrelated participants (490 male, 638 female) consisted of 872 adults and 256 children and adolescents (age *mean*=41.88 yrs, *SD*=21.56, *range* 0-72, see supplementary Figure S1 for density plot of age distribution). 95% of participants were born in Germany.

**Table 3.**
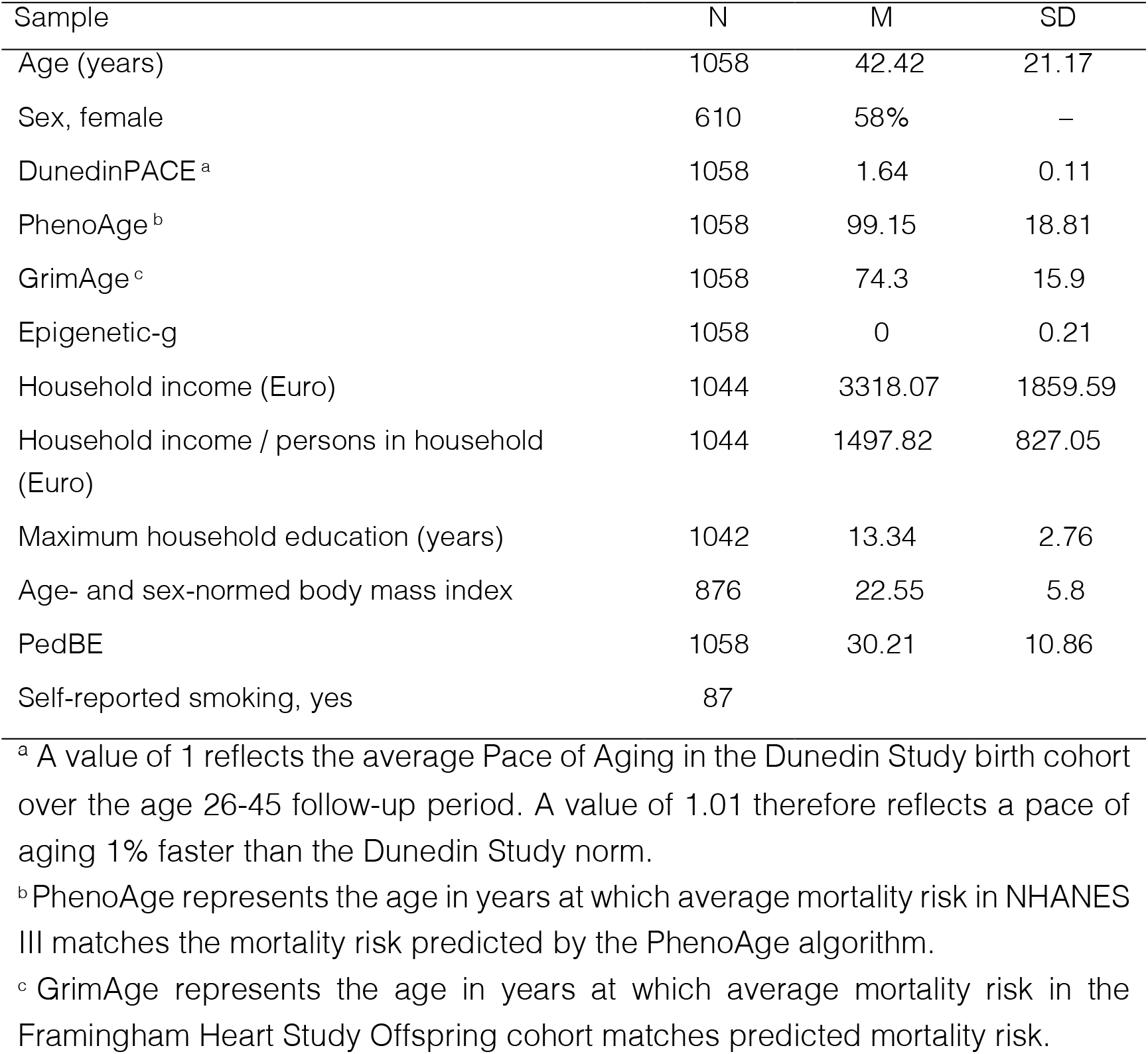
Descriptive statistics of the analytic sample after DNA-methylation based exclusions (N= 1058).

| Sample | N | M | SD |
| --- | --- | --- | --- |
| Age (years) | 1058 | 42.42 | 21.17 |
| Sex, female | 610 | 58% | – |
| DunedinPACE <sup>a</sup> | 1058 | 1.64 | 0.11 |
| PhenoAge <sup>b</sup> | 1058 | 99.15 | 18.81 |
| GrimAge <sup>c</sup> | 1058 | 74.3 | 15.9 |
| Epigenetic-g | 1058 | 0 | 0.21 |
| Household income (Euro) | 1044 | 3318.07 | 1859.59 |
| Household income / persons in household (Euro) | 1044 | 1497.82 | 827.05 |
| Maximum household education (years) | 1042 | 13.34 | 2.76 |
| Age- and sex-normed body mass index | 876 | 22.55 | 5.8 |
| PedBE | 1058 | 30.21 | 10.86 |
| Self-reported smoking, yes | 87 |  |  |

<sup>a</sup> A value of 1 reflects the average Pace of Aging in the Dunedin Study birth cohort over the age 26-45 follow-up period. A value of 1.01 therefore reflects a pace of aging 1% faster than the Dunedin Study norm.
<sup>b</sup> PhenoAge represents the age in years at which average mortality risk in NHANES III matches the mortality risk predicted by the PhenoAge algorithm.
<sup>c</sup> GrimAge represents the age in years at which average mortality risk in the Framingham Heart Study Offspring cohort matches predicted mortality risk.

## Measures

### DNA-methylation preprocessing and exclusions

DNA was extracted from buccal swabs collected using Isohelix IS SK-1S Dri-Capsules (Koellinger et al 2021). DNA extraction and methylation profiling was conducted by the Human Genomics Facility (HuGe-F) at the Erasmus Medical Center in Rotterdam, Netherlands. The Infinium MethylEPIC v1 manifest B5 kit (Illumina, Inc., San Diego, CA) was used to assess methylation levels at 865,918 CpG sites.

DNAm preprocessing was primarily conducted with Illumina’s GenomeStudio software and open-source *R* (version 4.2.0) packages ‘minfi’ (Aryee et al., 2014) and ‘ewastools’ (Heiss & Just, 2018). We generated 20 control metrics in GenomeStudio as described in the BeadArray

#### Controls Reporter Software Guide

from Illumina (note similar parameters can be computed using the ewastools ‘control_metrics()’ function). Samples falling below the Illumina-recommended cut-offs were flagged and further investigated. Flagged samples were classified as failed if 1. all types of poor bisulfite conversion and all types of poor bisulfite conversion background; 2. all types of bisulfite conversion background falling below 0.5; 3. all types of poor hybridization; 4. all types of poor specificity (excluded n=42).

As a second step, we identified unreliable data points resulting from low fluorescence intensities by filtering using detection p-values, calculated from comparing fluorescence intensities to a noise distribution. We removed probes with only background signal in a high proportion of samples (proportion of samples with detection p-value > 0.01 is > 0.1). We also removed probes for which a high proportion of samples had low bead numbers (proportion of samples with bead number < 3 is > 0.1). Further, we removed probes with SNPs at the CG or single base extension position as well as cross-reactive probes for EPIC arrays (McCartney et al., 2016; Pidsley et al., 2016).

We used minfi’s ‘preprocessNoob’ (Triche et al., 2013) to correct for background noise and color dye bias and ‘BMIQ’ to account for probe-type differences (Teschendorff & Widschwendter, 2012).

Cell composition was estimated using HEpiDISH, which is an iterative hierarchical version of the EpiDISH *R* package using robust partial correlations (https://github.com/sjczheng/EpiDISH). Because epithelial cell types are the dominant cell type in buccal samples, we applied a threshold of 0.5 for epithelial cell proportions to reliably call a “buccal sample” and excluded samples that failed this metric (n=28). All samples were from the same batch. Final analytic sample size after DNAm exclusions was N=1058. In GSE111165 blood samples, DNAm algorithms were residualized for reference-free cell composition and plate (Houseman et al., 2016).

### DNA-methylation algorithms

Our pre-registered analysis focused on two DNAm measures developed from blood DNAm data and which we had previously followed-up in saliva DNAm data (*i*.*e*., DunedinPACE and Epigenetic-*g)* as well as a buccal-based algorithm of chronological age to be used as a data quality control measure (*i*.*e*., PedBE). For comparative purposes, we report additional results for two further DNAm measures developed from blood DNAm, the PhenoAge and GrimAge clocks (Levine et al., 2018; Lu et al., 2019). We include these measures, which are among the best-evidenced DNAm biomarkers of aging, to help contextualize findings for DunedinPACE and Epigenetic-*g*. See Table 4 for description of DNA-methylation algorithm computations.

**Table 4.**
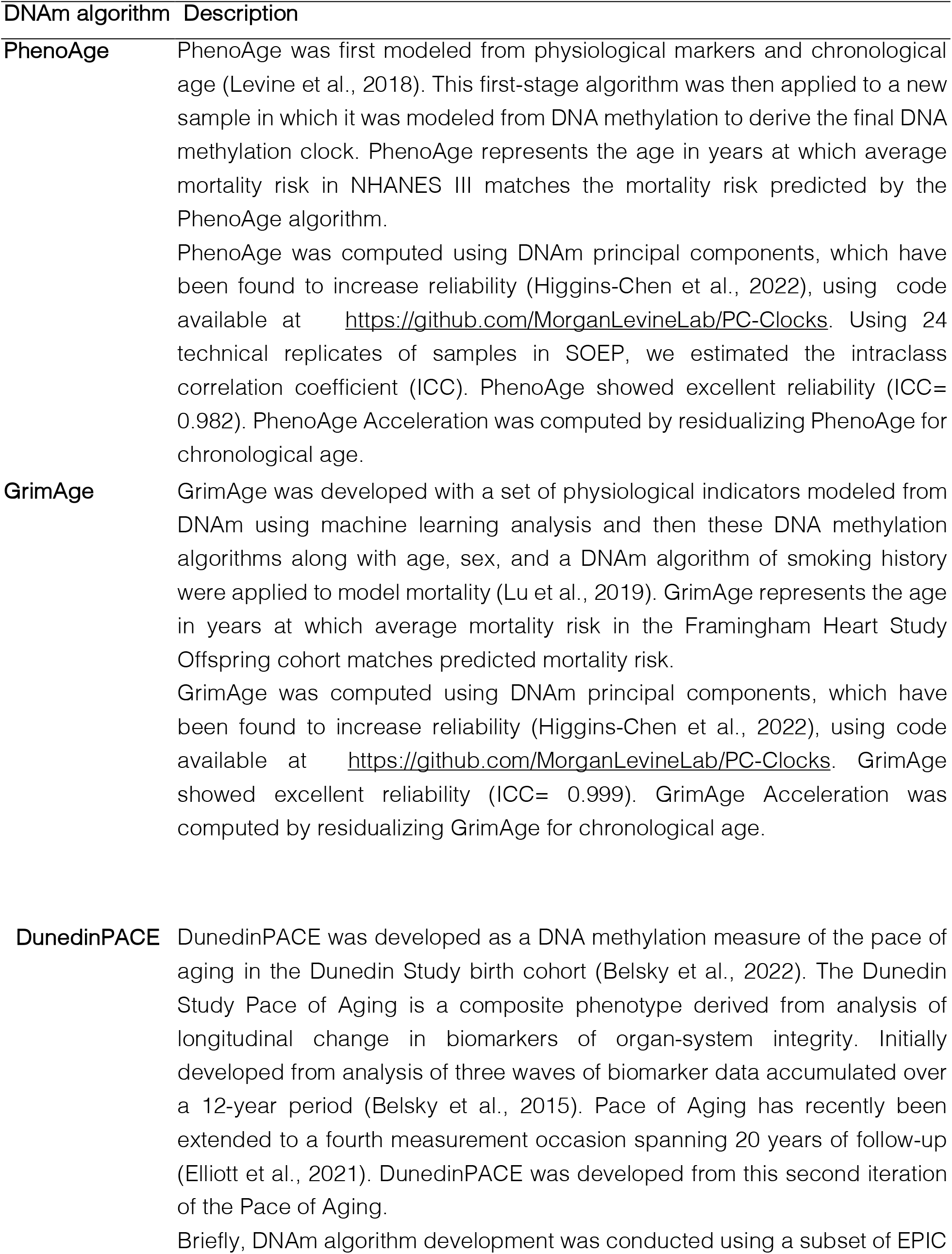
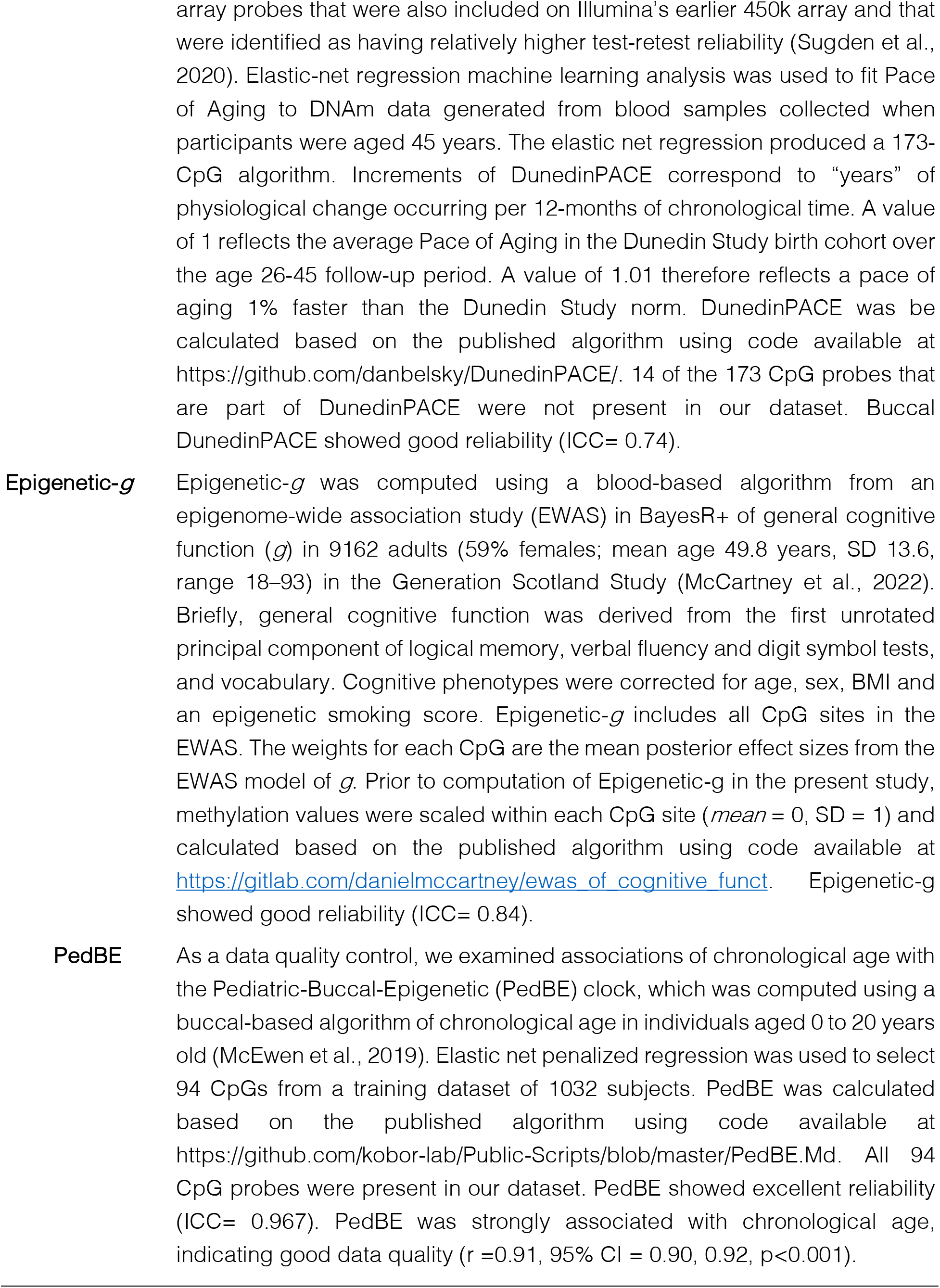
Description of DNA-methylation algorithm computations.

#### Socioeconomic status

We deviated from our pre-registered analysis plan by testing associations with socioeconomic status (average z-scored household income and education) rather than examining income and education separately, to reduce the number of statistical comparisons. Monthly household net income in Euros from all sources (e.g. employment, pensions, unemployment benefits, maternity benefits, higher education grants, military or civil service pay, compulsory child support, etc.) was reported by the self-defined head of household. In the 2% of cases with missing income values, information about determinants of household income and past data were used to impute estimated values (for more information see page 27 https://www.diw.de/documents/publikationen/73/diw_01.c.787445.de/diw_ssp0844.pdf)/. Household income was divided by the number of persons in the household and sqrt transformed to correct for skew (this deviated from our preregistration plan; sqrt-transformation improved normality of distribution more than log-transformation in shapiro wilks test).

Given the wide age range of participants, we indexed educational attainment as the highest degree obtained by any individual in the household. Educational attainment was converted to number of educational years (no degree = 7 years, lower school degree = 9 years, intermediary school = 10 years, degree for, a professional coll. = 12 years, high school degree = 13 years, other = 10 years) with additional occupational training added (apprenticeship = +1.5 years, technical schools (including health) = +2 years, civil servants apprenticeship = +1.5 years, higher technical college = +3 years, university degree = +5 years).

### Covariates

#### Body mass index (BMI)

Height (in cm) and weight (in kg) were measured via self-report and transformed to sex- and age-normed BMI z-scores

#### Smoking

Participant self-reported current or past smoking across multiple waves. Across questions and waves, if a participant ever responded that they smoked currently or in the past, they were identified as a smoker. If a participant ever responded that they never smoked and never responded that they did smoke, they were identified as a never-smoker.

## Declarations

### Ethics approval and consent to participate

Participants or the parents of minor participants consented to the archiving, extraction, and analysis of DNA-based measures. Ethical approval for DNA-based research was received by the Research Ethics Review Board of Vrije Universiteit Amsterdam, School of Business and Economics (application number 20181018.1.pkr730) and the Ethics Council of the Max Planck Society (application number 2019_16).

### Consent for publication

Not applicable.

### Availability of data and materials

The datasets used and/or analysed during the current study are available from the corresponding author on reasonable request.

### Authors’ contributions

Laurel Raffington developed the study concept and design, performed and supervised data analysis, and drafted the manuscript. Ted Schwaba performed data analysis and provided critical revisions. Muna Aikins provided critical revisions. David Richter and Gert G. Wagner developed the study concept and provided critical revisions. K. Paige Harden, Daniel W. Belsky, and Elliot M. Tucker-Drob developed the study concept and design, supervised data analysis, drafted the manuscript, and provided critical revisions. All authors approved the final manuscript as submitted and agree to be accountable for all aspects of the work.

## Acknowledgements

We thank the participants of SOEP-G.

## List of abbreviations

DNAm: DNA methylation
SES: Socioeconomic status

## Supplemental Results

### Supplemental Figures

**Supplementary Figure S1.**
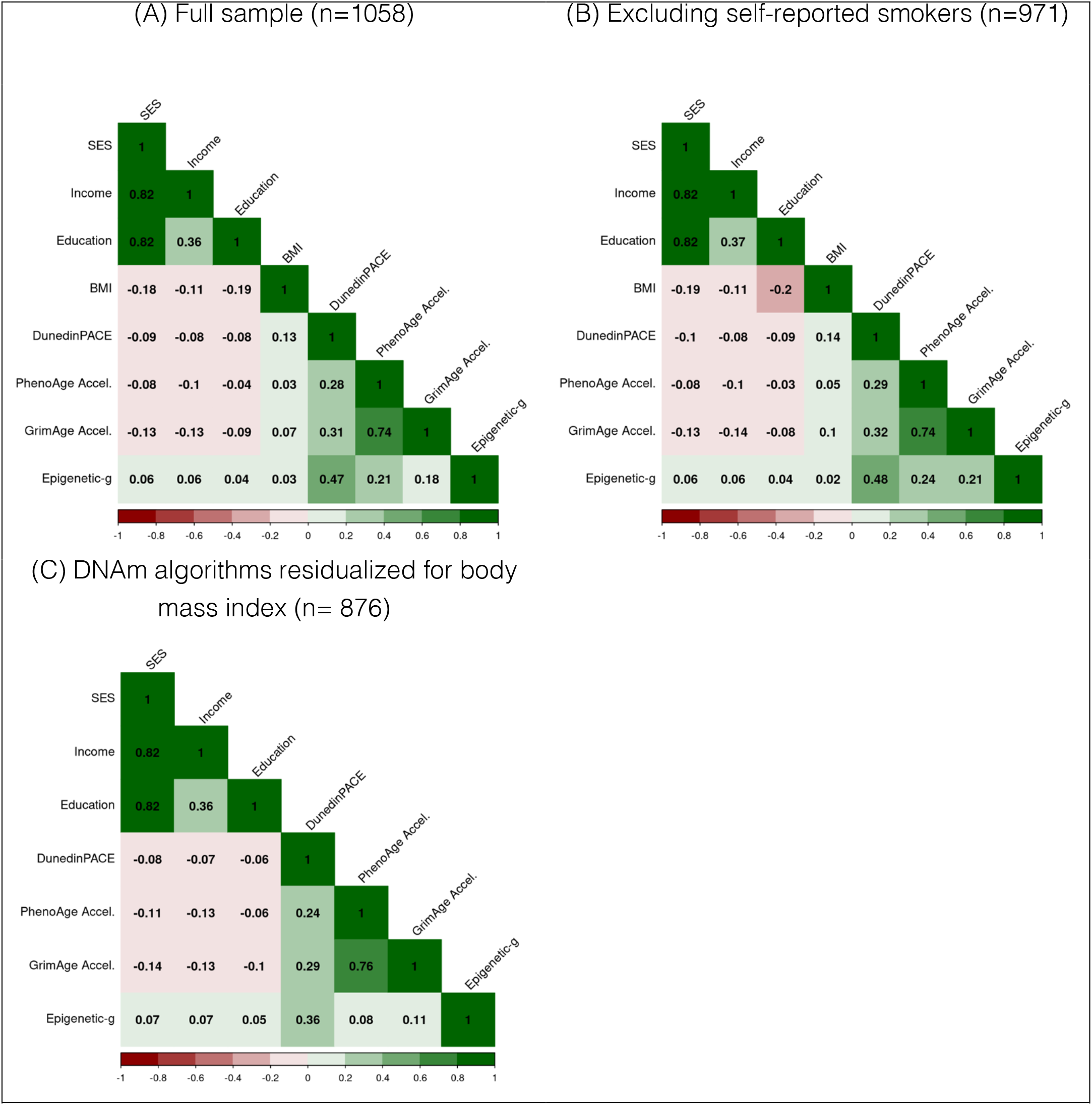
Correlation matrix of of socioeconomic variables with buccal DNA-methylation (DNAm) algorithms in (A) full sample, (B) excluding self-reported smokers, and (C) residualizing DNAm algorithms for body mass index (BMI). In all plots DNAm measures were residualized for chronological age. BMI was age- and sex-normed.

